# IFN-γ Orchestrates Coordinated Immunosuppression in Head and Neck Squamous Cell Carcinoma Through JAK-STAT-IRF8 Signaling: A Transcriptome-Wide Computational Analysis

**DOI:** 10.64898/2026.03.26.714228

**Authors:** Amir Mohamed Abdelhamid, Eslam E. Abd El-Fattah

## Abstract

**Background:** Interferon-gamma (IFN-γ) is the primary effector cytokine of adaptive anti-tumor immunity, yet it paradoxically induces a potent immunosuppressive tumor microenvironment (TME). The full mechanistic scope of this paradox in head and neck squamous cell carcinoma (HNSC) has not been characterized at the transcriptomic scale.

**Methods:** Using TCGA HNSC RNA-seq data (n = 522), we applied an integrated computational pipeline: Spearman correlation analysis, principal component analysis (PCA), UMAP, K-means clustering (k = 4), Random Forest regression, deep neural networks, permutation importance, JAK-STAT cascade mapping, and DNN-based transcriptome-wide mediation analysis across 57 IFN-γ pathway and 78 immunosuppressive genes.

**Results:** IFN-γ pathway activity was universally and positively correlated with six immunosuppressive axes, including checkpoints (CD274; LAG3; IDO1), Tregs, myeloid suppression, and tryptophan catabolism. K-means clustering identified four immunologically distinct tumor subgroups. DNN models predicted suppressive TME. Permutation importance identified IRF8 as the dominant mediator linking IFN-γ signaling to immunosuppression. DNN mediation analysis identified PDCD1LG2 (PD-L2) as the strongest intermediary between IFNG and PD-L1 regulation, followed by JAK2 and GBP5.

**Conclusions:** IFN-γ orchestrates coordinated immunosuppression in HNSC through JAK-STAT-IRF8 signaling. PDCD1LG2 and JAK2 are actionable mediators of this paradox, supporting combination strategies co-targeting IFN-γ-induced checkpoint induction and direct checkpoint blockade in HNSC immunotherapy.

**GRAPHICAL ABSTRACT:** 

## 1. INTRODUCTION

Head and neck squamous cell carcinoma (HNSC) represents the sixth most common malignancy worldwide, with over 900,000 new cases diagnosed annually. Despite multimodal treatment advances including surgery, radiotherapy, and chemotherapy, five-year survival rates remain below 50% for advanced-stage disease. The introduction of immune checkpoint inhibitors (ICIs) targeting the programmed death-1/programmed death-ligand 1 (PD-1/PD-L1) axis has provided clinical benefit in recurrent or metastatic HNSC, yet objective response rates with anti-PD-1 monotherapy remain modest at 13–18% in unselected patient populations [1–4]. This limited efficacy underscores the existence of potent immunosuppressive mechanisms within the HNSC tumor microenvironment (TME) that counteract antitumor immunity and necessitates deeper mechanistic understanding to guide rational combination immunotherapy strategies.

Interferon-gamma (IFN-γ), the sole member of the type II interferon family, constitutes a cardinal effector cytokine of adaptive antitumor immunity. Secreted principally by activated CD8^+^ cytotoxic T lymphocytes (CTLs) and natural killer (NK) cells upon tumor antigen recognition, IFN-γ binds the heterodimeric interferon-gamma receptor (IFNGR1/IFNGR2), initiating canonical Janus kinase 1/Janus kinase 2–signal transducer and activator of transcription 1 (JAK1/JAK2–STAT1) phosphorylation cascades. Activated STAT1 homodimers (gamma-activated factor, GAF) and STAT1–STAT2–interferon regulatory factor 9 (IRF9) heterotrimers (interferon-stimulated gene factor 3, ISGF3) translocate to the nucleus, binding gamma-activated sequences (GAS) and interferon-stimulated response elements (ISRE) to drive expression of interferon-stimulated genes (ISGs) [5–9]. These ISGs orchestrate a multifaceted antitumor program encompassing enhanced major histocompatibility complex class I (MHC-I) antigen presentation (human leukocyte antigen A/B/C [HLA-A/B/C], beta-2-microglobulin [B2M], transporter associated with antigen processing 1/2 [TAP1/TAP2]), recruitment of effector lymphocytes via CXC motif chemokine ligands (CXCL9, CXCL10, CXCL11), and direct antiproliferative and proapoptotic effects mediated by guanylate-binding proteins (GBP1–5) and interferon-induced proteins with tetratricopeptide repeats (IFIT1–3) [10–12].

Paradoxically, the same IFN-γ signaling axis that initiates antitumor immunity simultaneously drives expression of potent immunosuppressive mechanisms in a phenomenon termed adaptive immune resistance. CD274 (PD-L1) and PDCD1LG2 (PD-L2), encoding the primary ligands for the inhibitory receptor programmed cell death protein 1 (PD-1), are direct transcriptional targets of STAT1 and IRF1, providing a molecular explanation for the clinical observation that tumors with high IFN-γ pathway activity paradoxically display elevated checkpoint expression. Similarly, indoleamine 2,3-dioxygenase 1 (IDO1), the rate-limiting enzyme catalyzing tryptophan degradation to kynurenine—a metabolite that suppresses T-cell proliferation and promotes regulatory T-cell (Treg) differentiation—is robustly induced by IFN-γ via STAT1-dependent mechanisms. Beyond direct transcriptional regulation, IFN-γ orchestrates recruitment and polarization of immunosuppressive myeloid populations, including tumor-associated macrophages (TAMs) and myeloid-derived suppressor cells (MDSCs) through induction of colony-stimulating factor 1 (CSF1), C-C motif chemokine ligand 2 (CCL2), and related cytokines. IFN-γ further upregulates additional checkpoint molecules, including lymphocyte activation gene 3 (LAG3), T-cell immunoreceptor with Ig and ITIM domains (TIGIT), and hepatitis A virus cellular receptor 2 (HAVCR2, encoding TIM-3), creating a densely layered suppressive architecture [13–15].

This immunological duality—wherein IFN-γ serves as both the primary driver of antitumor immunity and the architect of feedback immunosuppression—carries profound therapeutic implications. Pretreatment IFN-γ gene signatures correlate with improved outcomes following ICI therapy across multiple cancer types, suggesting that baseline immune activation is necessary for response. Conversely, tumors displaying acquired resistance to PD-1 blockade frequently exhibit amplification of JAK-STAT pathway components and downstream immunosuppressive targets, including CD274, IDO1, and HAVCR2, indicating that excessive adaptive resistance can override checkpoint inhibition. The net clinical benefit of ICIs may therefore depend critically upon the quantitative balance between IFN-γ-driven tumor immunity and IFN-γ-induced immunosuppression. Rational design of combination immunotherapies targeting this balance requires comprehensive delineation of the molecular mediators linking IFN-γ pathway activation to specific suppressive mechanisms [16–18]. Despite extensive study of individual IFN-γ-induced immunosuppressive molecules, a systems-level characterization of the IFN-γ→immunosuppression regulatory architecture in HNSC remains absent. Prior investigations have largely focused on single pathway components or employed candidate gene approaches, leaving unresolved questions regarding: (1) which IFN-γ pathway genes exhibit strongest predictive power for distinct immunosuppressive axes; (2) whether HNSC tumors stratify into immunologically discrete subgroups based on coupled IFN-γ and suppressive TME profiles; (3) the relative contributions of JAK-STAT signaling components, interferon regulatory factors, and feedback regulators to immunosuppression prediction; and (4) the molecular intermediaries that mediate IFN-γ effects on specific checkpoint molecules such as PD-L1. The Cancer Genome Atlas (TCGA) HNSC dataset, comprising RNA-sequencing profiles from 522 primary tumors with comprehensive transcriptomic coverage, provides an unprecedented resource for genome-wide dissection of these questions.

In this study, we applied an integrated machine learning and deep learning framework to TCGA HNSC transcriptomic data to comprehensively map the IFN-γ immunosuppressive landscape. We employed random forest feature importance analysis and deep neural network (DNN) permutation importance testing to identify master regulators of immunosuppression prediction from 57 IFN-γ pathway genes. Unsupervised clustering based on joint IFN-γ pathway and suppressive TME scores revealed four immunologically distinct HNSC subgroups, including an immune-hot, maximally suppressed phenotype of primary therapeutic interest. DNN models achieved R^2^ = 0.65–0.85 across six suppressive categories (checkpoint molecules, Treg-induction genes, anti-inflammatory cytokines, tryptophan-kynurenine metabolism, STAT3-dependent suppression, and MDSCs/TAMs). Critically, permutation importance analysis identified interferon regulatory factor 8 (IRF8) as the dominant predictor of immunosuppression by an extraordinary margin (ΔR^2^ = 0.67), exceeding the second-ranked gene by more than 15-fold. Finally, transcriptome-wide DNN-based mediation analysis revealed that PDCD1LG2 (PD-L2) acts as the primary molecular intermediary linking IFNG expression to CD274 (PD-L1) upregulation, improving predictive accuracy by 50% (ΔR^2^ = +0.212) over IFNG alone. Collectively, these findings delineate the mechanistic architecture of adaptive immune resistance in HNSC, identify IRF8 as a master transcriptional integrator of IFN-γ-mediated immunosuppression, and nominate rational therapeutic targets for combinatorial immune checkpoint blockade strategies.

## 2. MATERIALS AND METHODS

### 2.1 Data Source and Preprocessing

Gene expression data for head and neck squamous cell carcinoma (HNSC) were obtained from The Cancer Genome Atlas (TCGA) program via the National Cancer Institute (NCI) Genomic Data Commons (GDC) data portal (https://portal.gdc.cancer.gov/). RNA-sequencing (RNA-seq) data were downloaded as gene-level read count matrices and imported using the pandas library (version 1.5.3) in Python 3.10. Quality control filtering excluded samples with fewer than 10 detected genes (minimum sample threshold = 10). Gene-level count matrices were filtered to retain only uniquely named genes; duplicate gene identifiers were resolved by retaining the first occurrence after removing gene_id and gene_type metadata columns derived from GENCODE annotation. To stabilize variance and approximate normal distributions, log_2_(x + 1) transformation was applied to all raw count values, where x represents the read count for each gene. Missing expression values within each gene row were imputed using row-wise median imputation to preserve gene-specific expression distributions while minimizing bias from absent data. TCGA sample barcodes were harmonized by truncating to the first 12 characters of standard TCGA identifiers to ensure consistent sample mapping across data matrices. The final curated HNSC dataset comprised 522 primary tumor samples with complete transcriptomic profiles. All computational analyses were performed in Python 3.10 using NumPy (version 1.24.2) for numerical operations, pandas (version 1.5.3) for data manipulation, SciPy (version 1.10.1) for statistical testing, scikit-learn (version 1.2.2) for machine learning algorithms, matplotlib (version 3.7.1) and seaborn (version 0.12.2) for data visualization, and NetworkX (version 3.1) for network analysis [19–21].

### 2.2 Literature-Curated Gene Sets

Two comprehensive, literature-curated gene-set dictionaries formed the molecular foundation of this analysis. The IFN-γ Pathway Gene Set comprised 57 genes organized into four functional subcategories based on established roles in interferon-gamma signaling biology. The *IFNG_core* subcategory (n = 3 genes) included the ligand and receptor components: *IFNG* (interferon-gamma), *IFNGR1* (interferon-gamma receptor 1), and *IFNGR2* (interferon-gamma receptor 2). The *JAK_STAT* subcategory (n = 16 genes) encompassed proximal signaling mediators: *JAK1* (Janus kinase 1), *JAK2* (Janus kinase 2), *TYK2* (tyrosine kinase 2), *STAT1*–*STAT3* (signal transducer and activator of transcription 1–3), *IRF1*–*IRF3* and *IRF7*–*IRF9* (interferon regulatory factors 1–3 and 7–9), *SOCS1* and *SOCS3* (suppressor of cytokine signaling 1 and 3), and *PIAS1* and *PIAS3* (protein inhibitor of activated STAT 1 and 3).

The *IFNG_targets* subcategory (n = 28 genes) comprised canonical interferon-stimulated genes (ISGs) including guanylate-binding proteins (*GBP1*–*GBP5*), CXC motif chemokine ligands (*CXCL9*, *CXCL10*, *CXCL11*), major histocompatibility complex class I molecules (*HLA-A*, *HLA-B*, *HLA-C*), MHC class II components (*HLA-DRA*, *HLA-DRB1*), antigen processing machinery (*B2M* [beta-2-microglobulin], *TAP1* and *TAP2* [transporter associated with antigen processing 1 and 2], *TAPBP* [TAP-binding protein]), antiviral effectors (*MX1* [MX dynamin like GTPase 1], *OAS1* and *OAS2* [2’-5’-oligoadenylate synthetase 1 and 2], *IFIT1*–*IFIT3* [interferon-induced protein with tetratricopeptide repeats 1–3]), and ISGylation pathway components (*ISG15*, *ISG20*, *HERC5* [HECT and RLD domain containing E3 ubiquitin protein ligase 5], *USP18* [ubiquitin specific peptidase 18]).

The *IFNG_feedback* subcategory (n = 14 genes) included negative regulators and pathway modulators: *TRIM63* (tripartite motif containing 63), ubiquitin components (*UBB*, *UBC*), nuclear factor kappa B (NF-κB) pathway genes (*NFKB1*, *NFKB2*, *RELA*, *RELB*, *REL*), innate immune signaling kinases (*IKBKE* [inhibitor of nuclear factor kappa B kinase subunit epsilon], *TBK1* [TANK binding kinase 1]), and cytosolic DNA sensing pathway components (*STING1* [stimulator of interferon response cGAMP interactor 1], *CGAS* [cyclic GMP-AMP synthase], *TRIM56*, *TRIM32*).

The Immunosuppressive Gene Set encompassed 78 genes organized into six mechanistically distinct functional axes. The *Checkpoints_induced* axis (n = 15 genes) comprised canonical and emerging immune checkpoint molecules: *CD274* (programmed death-ligand 1 [PD-L1]), *PDCD1LG2* (programmed death-ligand 2 [PD-L2]), *CD80* and *CD86* (co-stimulatory molecules B7-1 and B7-2), *HAVCR2* (hepatitis A virus cellular receptor 2, encoding TIM-3), *LAG3* (lymphocyte activation gene 3), *TIGIT* (T cell immunoreceptor with Ig and ITIM domains), *IDO1* and *IDO2* (indoleamine 2,3-dioxygenase 1 and 2), *CTLA4* (cytotoxic T-lymphocyte associated protein 4), *VSIR* (V-set immunoregulatory receptor, encoding VISTA), sialic acid-binding immunoglobulin-like lectins (*SIGLEC7*, *SIGLEC9*), and the CD47-SIRPα don’t-eat-me axis (*CD47*, *SIRPA* [signal regulatory protein alpha]).

The *Treg_induction* axis (n = 8 genes) included regulatory T-cell master regulators and functional markers: *FOXP3* (forkhead box P3), *IL2RA* (interleukin-2 receptor alpha chain, CD25), *IKZF2* (IKAROS family zinc finger 2, encoding Helios), *TNFRSF18* (tumor necrosis factor receptor superfamily member 18, encoding GITR), adenosine-generating ectonucleotidases (*ENTPD1* [ectonucleoside triphosphate diphosphohydrolase 1, CD39], *NT5E* [5’-nucleotidase ecto, CD73]), *CCR8* (C-C motif chemokine receptor 8), and *TIGIT*.

The *Anti_inflam_cyto* axis (n = 10 genes) encompassed immunosuppressive cytokines and growth factors: *IL10* (interleukin-10), transforming growth factor beta isoforms (*TGFB1*, *TGFB2*, *TGFB3*), *IL6* (interleukin-6), regulatory interleukins (*IL27*, *IL35*, *IL37*), and vascular endothelial growth factors (*VEGFA*, *VEGFB*).

The *MDSCs_TAMs* axis (n = 15 genes) comprised myeloid-derived suppressor cell (MDSC) and tumor-associated macrophage (TAM) functional markers: *ARG1* and *ARG2* (arginase 1 and 2), *NOS2* (nitric oxide synthase 2), damage-associated molecular patterns (*S100A8*, *S100A9*), myeloid lineage markers (*CD33*, *ITGAM* [integrin subunit alpha M, CD11b], *CD68*), M2 macrophage-associated receptors (*MRC1* [mannose receptor C-type 1, CD206], *CD163*, *MSR1* [macrophage scavenger receptor 1]), colony-stimulating factor signaling (*CSF1R* [colony stimulating factor 1 receptor], *CSF1*), and Th2-polarizing cytokines (*IL4*, *IL13*).

The *Tryptophan_kynurenine* axis (n = 9 genes) included enzymes of the tryptophan-kynurenine metabolic pathway: *IDO1* and *IDO2*, kynurenine pathway enzymes (*KYNU* [kynureninase], *HAAO* [3-hydroxyanthranilate 3,4-dioxygenase], *ACMSD* [aminocarboxymuconate semialdehyde decarboxylase], *AFMID* [arylformamidase], *KMO* [kynurenine 3-monooxygenase]), the alternative tryptophan-degrading enzyme *TDO2* (tryptophan 2,3-dioxygenase), and the tryptophan-tRNA ligase *WARS1* (tryptophanyl-tRNA synthetase 1).

The *STAT3_suppress* axis (n = 10 genes) encompassed STAT3-dependent oncogenic and immunosuppressive programs: *STAT3* itself, upstream activators (*IL6*, *IL10*), anti-apoptotic targets (*BCL2* [BCL2 apoptosis regulator], *MCL1* [MCL1 apoptosis regulator, BCL2 family member]), angiogenic factors (*VEGFA*), oncogenic drivers (*MYC* [MYC proto-oncogene], *HIF1A* [hypoxia inducible factor 1 subunit alpha]), and epithelial-mesenchymal transition regulators (*TWIST1*, *SNAI1* [snail family transcriptional repressor 1]). All gene set annotations were curated from primary literature sources, including landmark studies defining IFN-γ signaling architecture and immune checkpoint biology.

### 2.3 Composite Scoring, Correlation Analysis, and Dimensionality Reduction

Composite IFN-γ pathway and suppressive tumor microenvironment (TME) scores were computed as arithmetic means of log_2_(x + 1)-transformed expression values across all detected member genes within each respective gene set, providing continuous quantitative metrics of pathway activity at the single-tumor level. Samples were dichotomized into IFNG-High and IFNG-Low groups by median split of the composite IFN-γ score to facilitate categorical comparisons of gene expression profiles. Pairwise Spearman rank correlation coefficients (ρ) between all IFN-γ pathway genes and all immunosuppressive genes were computed using SciPy’s spearmanr function, with statistical significance assessed at a Bonferroni-corrected threshold of p < 0.05 to control family-wise error rate across the large number of pairwise comparisons. Principal component analysis (PCA) was applied to a standardized (z-score normalized) combined feature matrix comprising all 57 IFN-γ pathway genes and 78 immunosuppressive genes using scikit-learn’s StandardScaler for normalization and PCA for dimensionality reduction. Uniform Manifold Approximation and Projection (UMAP), a non-linear dimensionality reduction technique that preserves both local and global data structure, was applied to the same standardized combined feature matrix using the umap-learn Python package with parameters n_neighbors = 15 (controlling local neighborhood size) and min_dist = 0.1 (governing minimum separation between embedded points) [21–25].

### 2.4 Unsupervised Clustering

K-means clustering (k = 4 clusters, n_init = 10 initializations, random_state = 42 for reproducibility) was applied to PCA-reduced feature representations using scikit-learn’s KMeans implementation to partition the HNSC cohort into immunologically distinct molecular subgroups. The choice of k = 4 was informed by silhouette score analysis and biological interpretability. Cluster-level gene expression profiles were computed as z-score-normalized mean expression values for each gene across samples within each cluster and visualized as heatmaps with hierarchical clustering of genes using Euclidean distance and Ward linkage [26].

### 2.5 Random Forest and Deep Neural Network Modelling

Separate Random Forest (RF) regression models (n_estimators = 200 decision trees, random_state = 42) were trained to predict each of the six suppressive TME group scores from the 57 IFN-γ pathway gene expression profiles using an 80/20 train-test split for model evaluation. Random Forest feature importance was quantified as the mean decrease in node impurity (Gini importance) averaged across all decision trees in the ensemble, providing a measure of each gene’s contribution to predictive accuracy. Multi-output deep neural network (DNN) models were constructed using scikit-learn’s MLPRegressor (multilayer perceptron regressor) with a four-hidden-layer architecture (128–64–32–16 neurons), rectified linear unit (ReLU) activation functions to introduce non-linearity, L_2_ regularization (alpha = 0.01) to prevent overfitting, and the Adam (adaptive moment estimation) optimizer for gradient-based parameter updating. DNN models were trained simultaneously on all six suppressive group scores as a multi-output regression task, with model performance assessed by coefficient of determination (R^2^) on held-out test data, where R^2^ = 1 − (sum of squared residuals / total sum of squares) represents the proportion of variance explained by the model.

### 2.6 Permutation Importance, Network Analysis, and JAK-STAT Cascade Mapping

Permutation importance, a model-agnostic technique for assessing feature importance by measuring the decrease in model performance when individual features are randomly shuffled, was computed on the fitted DNN using scikit-learn’s permutation_importance function with 10 independent repetitions per gene. The mean decrease in R^2^ (ΔR^2^) upon permutation of each gene’s expression values was used as the primary metric for ranking gene importance, with larger ΔR^2^ values indicating greater predictive contribution. A gene co-expression network was constructed to visualize functional relationships between IFN-γ pathway genes and immunosuppressive genes, with network edges drawn between gene pairs exhibiting |Spearman ρ| > 0.35 and Bonferroni-corrected p < 0.01, ensuring statistical rigor while focusing on moderate-to-strong correlations. Network visualization employed NetworkX’s spring layout algorithm (Fruchterman-Reingold force-directed layout) to position nodes such that highly correlated genes cluster spatially. A dedicated JAK-STAT signaling cascade heatmap was constructed by organizing pathway genes according to their hierarchical position in the canonical signaling flow (Receptor → JAK → STAT → IRF → Targets → Feedback) and displaying their expression patterns across samples sorted by ascending composite IFN-γ pathway score.

### 2.7 DNN-Based Mediation Analysis

To systematically identify molecular intermediaries linking *IFNG* expression to *CD274* (PD-L1) upregulation, we implemented a DNN-based mediation analysis framework. A baseline single-input DNN model (three hidden layers: 64–32–16 neurons; ReLU activation; Adam optimizer) was trained using *IFNG* expression alone as the predictor of *CD274* expression, achieving a baseline R^2^ = 0.426 on test data. For each candidate mediator gene across the entire transcriptome (n ≈ 20,000 protein-coding genes), a two-input DNN model (*IFNG* + candidate mediator gene) with identical architecture was trained to predict *CD274* expression. The improvement in predictive accuracy conferred by adding each candidate mediator was quantified as ΔR^2^ = R^2^_mediator_ − R^2^_baseline_, where positive ΔR^2^ values indicate that the mediator gene carries additional information beyond *IFNG* alone for predicting PD-L1 expression.

To complement the DNN-based ΔR^2^ metric, an indirect effect score was computed for each candidate mediator as the product of two Spearman correlation coefficients: r(*IFNG*, mediator gene) × r(mediator gene, *CD274*). This indirect score provides an estimate of the degree to which a mediator gene lies on a correlation path connecting *IFNG* to *CD274*, with high values indicating strong bidirectional correlations consistent with a mediation role. Top mediator genes were ranked by ΔR ^2^ and visualized through multiple complementary representations including: (1) horizontal bar plots showing the top 30 mediators ranked by ΔR^2^; (2) two-dimensional bubble plots with indirect score on the x-axis, ΔR^2^ on the y-axis, and bubble size proportional to the Spearman correlation between mediator and *CD274*; (3) DNN architecture diagrams illustrating the five-mediator model structure (*IFNG* + top 5 mediators → hidden layers → *CD274* output); and (4) expression heatmaps displaying the top 25 mediator genes across all HNSC samples sorted by *IFNG* expression to visualize co-regulation patterns. This comprehensive mediation analysis framework enabled systematic, genome-wide identification of the molecular intermediaries most critical for transducing IFN-γ signaling into PD-L1 expression [27]. The methodology working pipeline is illustrated in **Figure 2.3.**

## RESULTS

### 3.1 IFN-γ Pathway Expression Landscape and JAK-STAT Cascade in HNSC

To characterize the baseline IFN-γ signaling landscape, we generated a comprehensive heatmap of 57 IFN-γ pathway genes across 522 HNSC samples sorted by descending IFNG expression (Figure 3A). The heatmap revealed a graded, coordinated expression continuum: IFNG-high samples (left) showed broad co-upregulation of JAK-STAT components (STAT1, STAT2, IRF1, IRF7), IFN-γ target genes (GBP1–5, CXCL9–11, HLA-A/B/C, B2M, TAP1), and feedback regulators. STAT1 showed the highest dynamic range among JAK-STAT genes. B2M and classical HLA molecules maintained constitutively high expression across most samples, consistent with ubiquitous antigen presentation. TRIM63 displayed a striking inverse pattern—high in IFNG-low samples—consistent with a negative regulatory role attenuating IFN-γ responses.

**Figure 1.**
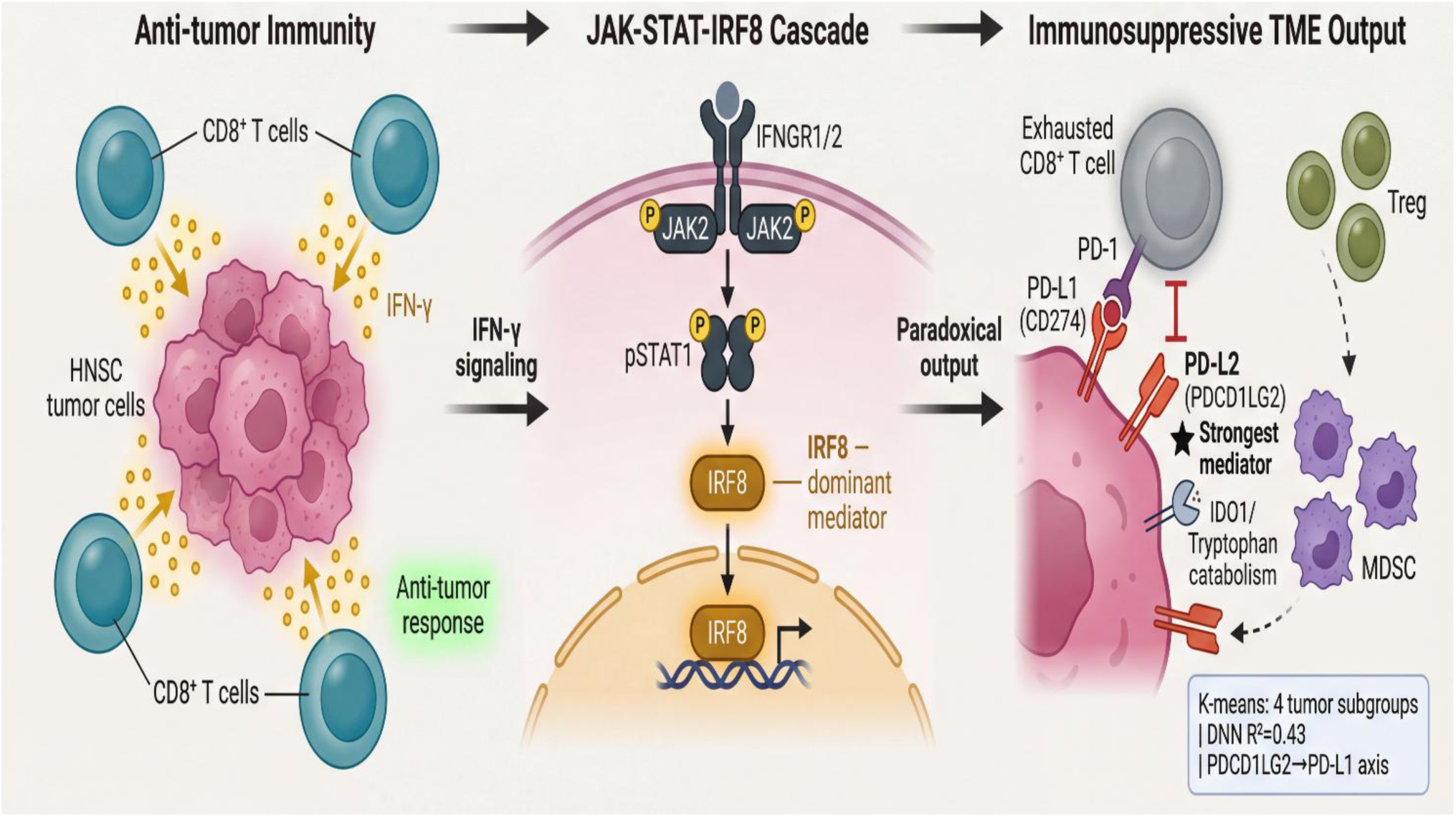
IFN-γ/JAK-STAT-IRF8 signaling drives paradoxical immunosuppression in HNSCC. CD8⁺ T cell-derived IFN-γ activates JAK2/pSTAT1 signaling in tumor cells, inducing IRF8 as the dominant downstream mediator. Rather than sustaining anti-tumor immunity, this cascade paradoxically upregulates PD-L1/PD-L2, IDO1/tryptophan catabolism, MDSCs, and Tregs, driving T cell exhaustion.

**Figure 2.**
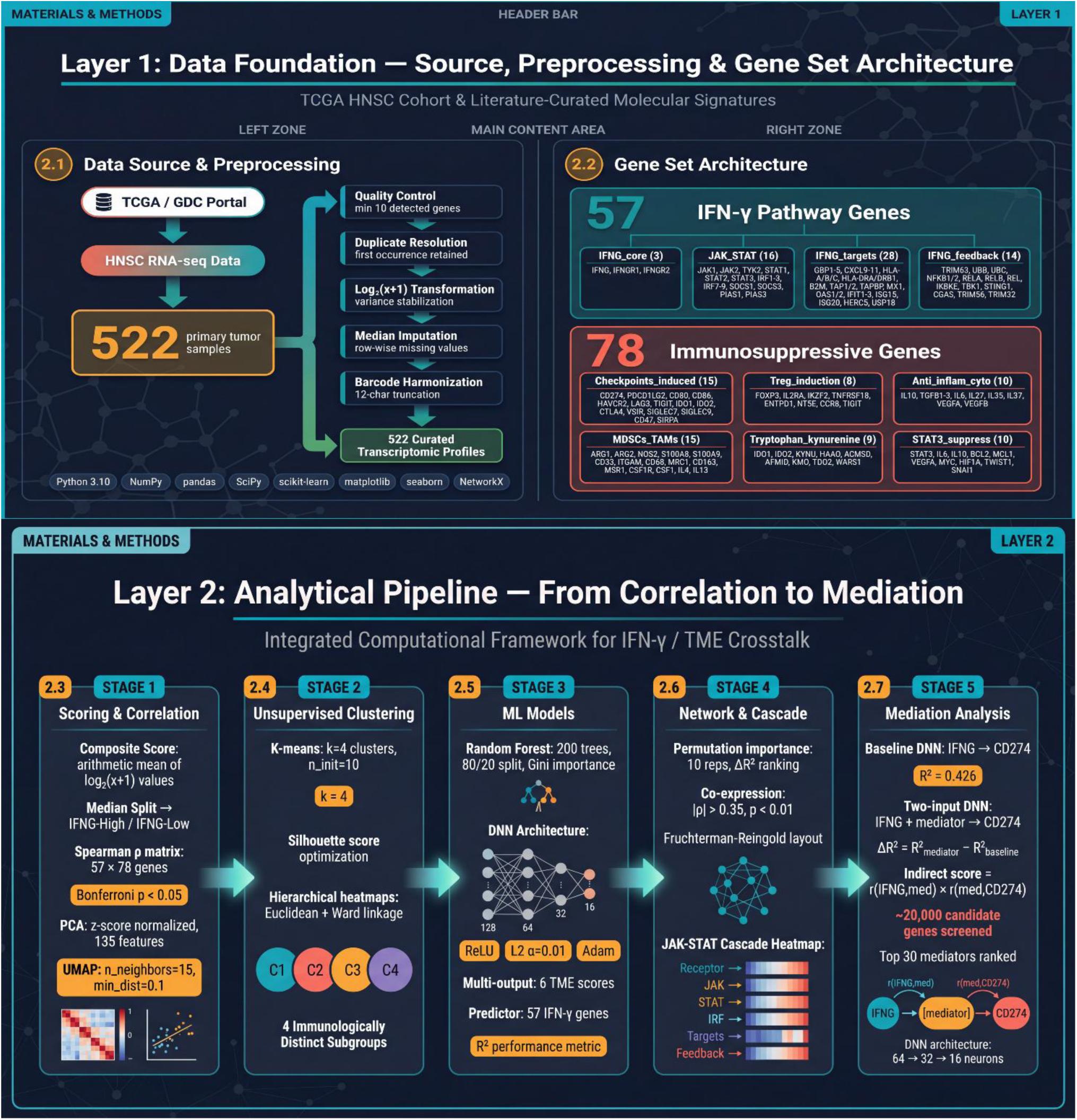
Integrated computational framework for analyzing IFN-γ/TME crosstalk in HNSCC. TCGA HNSC RNA-seq data (n=522) were preprocessed and analyzed using a curated 135-gene set comprising 57 IFN-γ pathway genes and 78 immunosuppressive genes across six functional categories. The pipeline integrated K-means clustering (k=4), Random Forest, and deep neural network (DNN: 128→64→32→16, R²=0.426) models alongside JAK-STAT cascade network analysis with permutation importance and co-expression thresholding (|ρ|>0.35). Mediation analysis screening ∼20,000 candidate genes identified top mediators of the IFNG→CD274 axis quantified by indirect scores r(IFNG,med)×r(med,CD274), implemented in Python using scikit-learn, NetworkX, and associated libraries.

**Figure 3.**
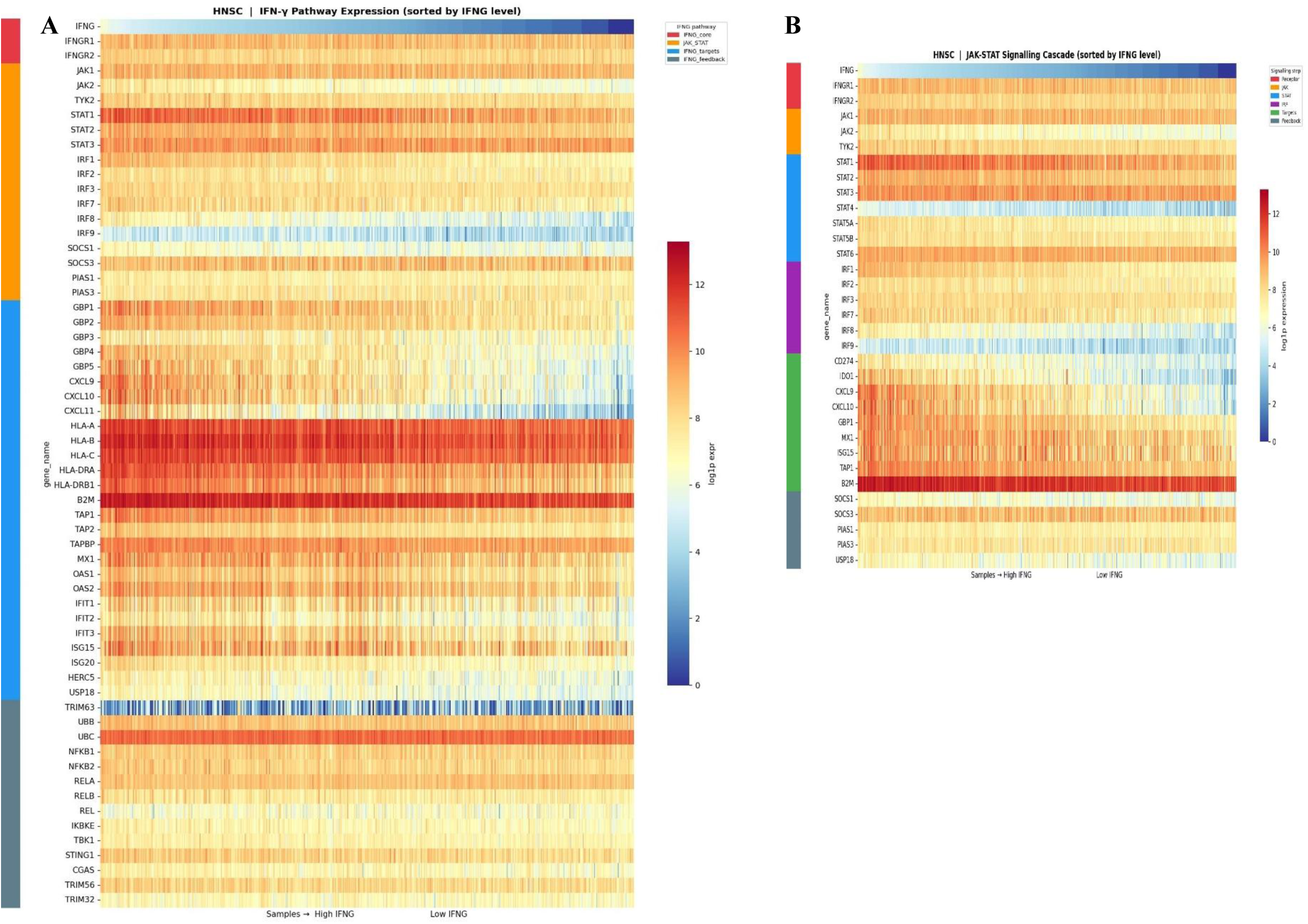
IFN-γ Pathway Expression Landscape and JAK-STAT Signaling Cascade in HNSC. (A) Heatmap of log1p-transformed expression for all 57 IFN-γ pathway genes across TCGA HNSC samples (n = 522) sorted left-to-right by descending IFNG expression. Row color annotations (left sidebar) denote subgroup membership: IFNG_core (red), JAK_STAT (orange), IFNG_targets (blue), IFNG_feedback (grey). (B) Dedicated JAK-STAT signaling cascade heatmap with genes organized by signaling step: Receptor (red), JAK kinases (orange), STAT transcription factors (blue), IRF family members (purple), ISG targets (green), and feedback regulators (grey). Samples are sorted by composite IFN-γ score (high → low, left to right). Colour scale in both panels: blue (low) to red (high) log1p expression. STAT1 shows the highest dynamic range among JAK-STAT genes; IRF8 displays a bimodal distribution; TRIM63 shows an inverse expression pattern.

The dedicated JAK-STAT cascade heatmap (Figure 3B) confirmed STAT1 as the dominant STAT isoform activated in IFNG-high tumors, with substantially higher dynamic range than STAT2, STAT3, or other STAT family members. Among IRF family members, IRF1 and IRF7 showed strong IFNG-correlated expression, while IRF8 displayed a bimodal distribution—intensely expressed in a subset of IFNG-high tumors but near-absent in others—presaging its identification as the dominant permutation importance feature. IRF9 was broadly low across the cohort. Key ISG targets (CD274, IDO1, CXCL9/10, GBP1, MX1, ISG15, TAP1, B2M) showed graded upregulation with increasing IFNG score. Feedback regulators (SOCS1, SOCS3, PIAS1/3, USP18) were co-upregulated in IFNG-high tumors, consistent with negative feedback on JAK-STAT signaling.

### 3.2 Genome-Wide Correlations Between IFN-γ Pathway and Immunosuppressive Axes

Pairwise Spearman correlations between all IFN-γ pathway genes and immunosuppressive genes (Bonferroni p < 0.05) revealed a striking predominance of positive correlations, with JAK-STAT genes (STAT1, IRF1, IRF7) showing the strongest positive associations with checkpoint-induced (CD274, LAG3, IDO1) and Treg-induction (FOXP3, CCR8) genes (Figure 4A). Negative correlations were sparse, concentrated in the ARG1 and VEGFA region of the MDSCs_TAMs axis, suggesting partial decoupling of myeloid suppression from canonical IFN-γ signaling.

**Figure 4.**
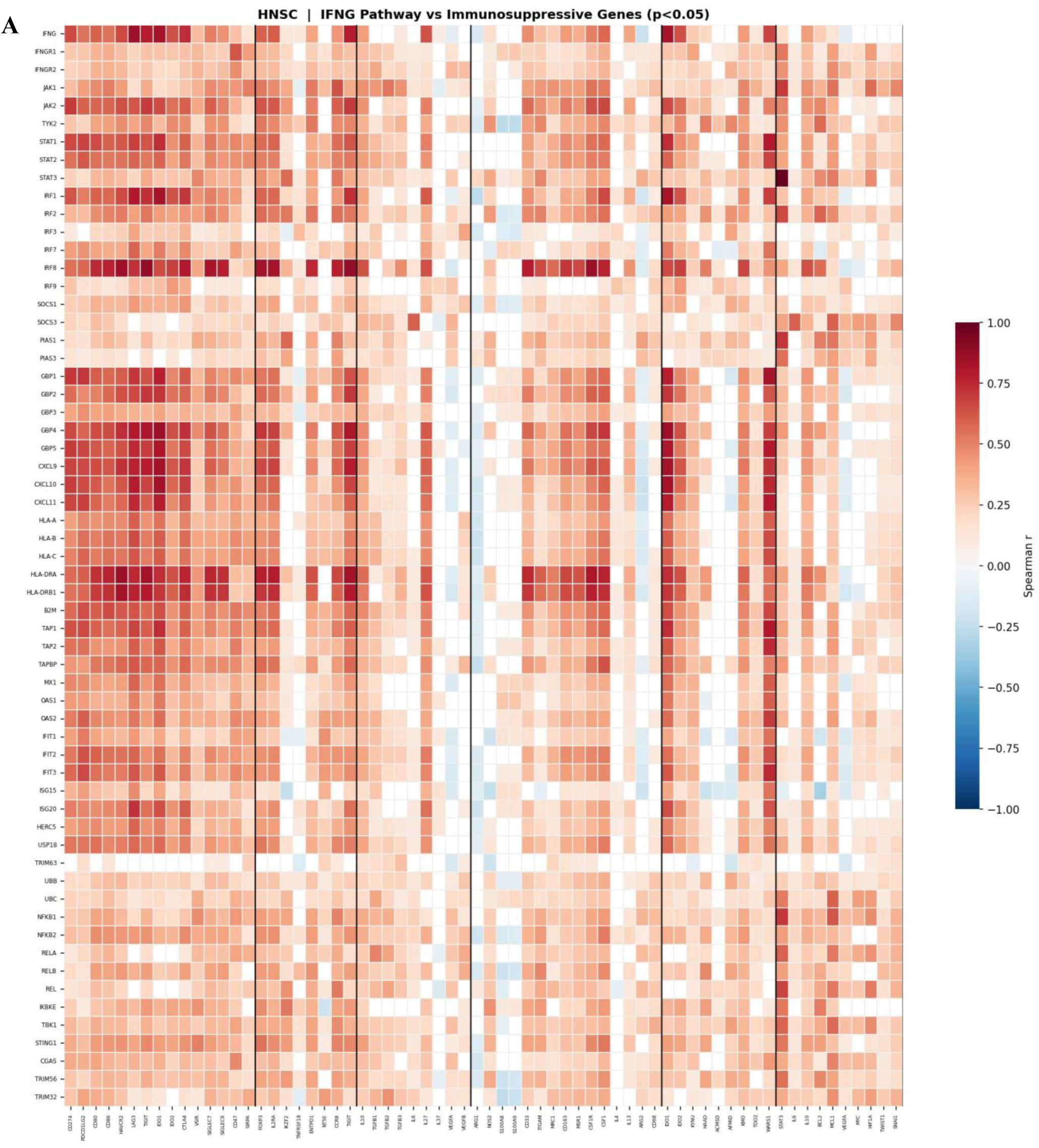
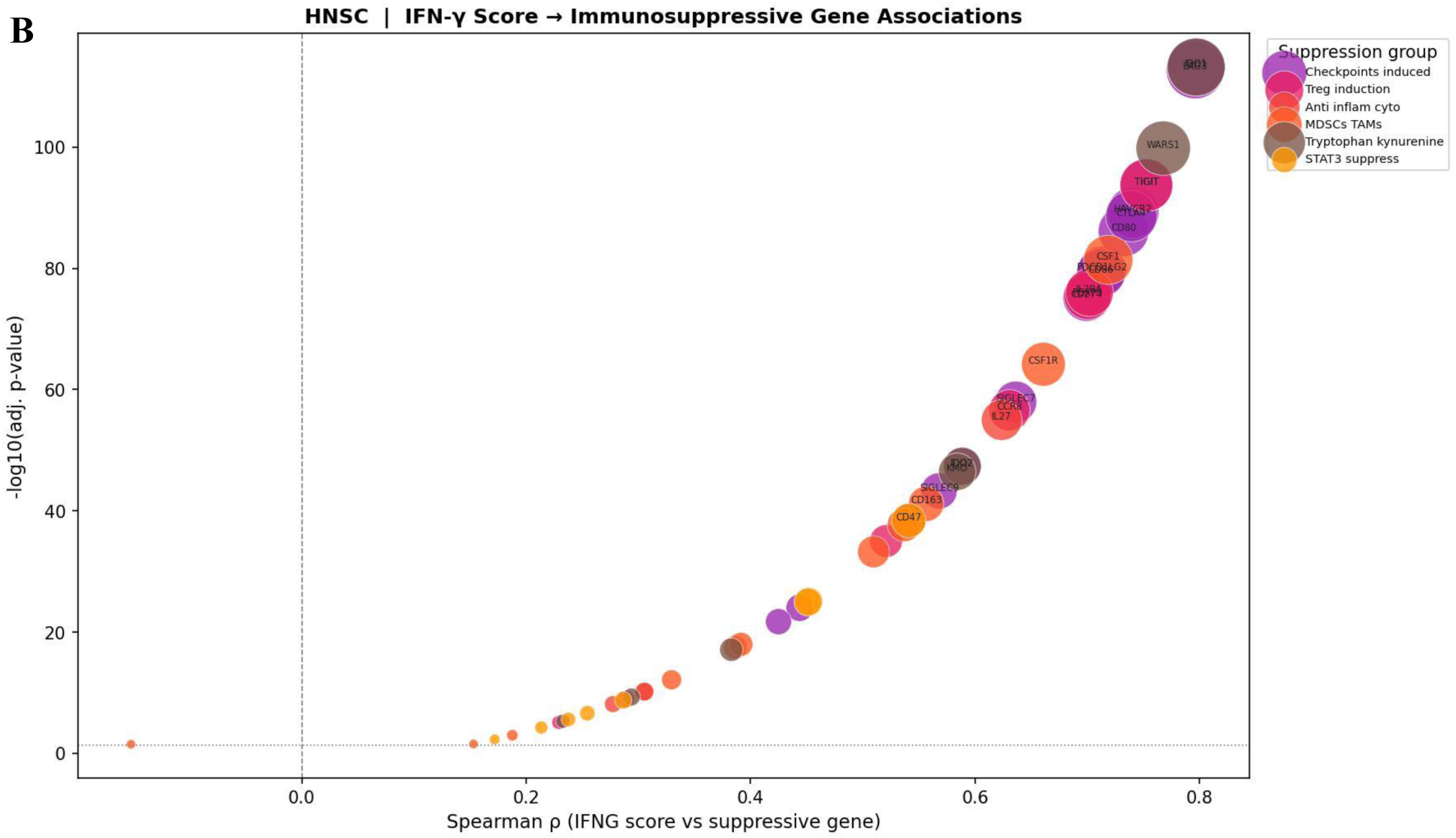
Genome-Wide Correlations Between IFN-γ Pathway and Immunosuppressive Gene Axes. (A) Spearman ρ heatmap between IFN-γ pathway genes (rows) and immunosuppressive genes (columns) in HNSC (Bonferroni p < 0.05). Non-significant pairs are masked (white). Vertical black lines demarcate the six immunosuppressive gene groups (labelled at the bottom, color-coded by group). Color scale: blue (ρ = −1) to red (ρ = +1). The predominance of red (positive correlations) indicates that IFN-γ pathway activation broadly co-occurs with immunosuppressive gene upregulation across all six axes. (B) Bubble plot of Spearman ρ between the composite IFN-γ pathway score and each immunosuppressive gene (x-axis) versus −log10(Bonferroni-adjusted p-value) (y-axis). Bubble size is proportional to −log10(padj). Bubble color indicates the immunosuppressive gene category. The horizontal dotted line denotes the p = 0.05 significance threshold. IDO1 and LAG3 (ρ = 0.80) are the strongest correlates of the IFN-γ score, while ARG1 (ρ = −0.15) is the sole negatively correlated gene.

The IFN-γ score bubble plot (Figure 4B) quantified associations between the composite IFN-γ pathway score and each immunosuppressive gene. All significant associations were positive (Spearman ρ range: 0.10–0.82). The strongest associations were IDO1 (ρ = 0.80, p_adj_ = 9.1×10^−116^), LAG3 (ρ = 0.80, p_adj_ = 2.6×10^−115^), TIGIT (ρ = 0.75, p_adj_ = 2.8×10^−96^), HAVCR2/TIM-3 (ρ = 0.74, p_adj_ = 8.8×10^−92^), FOXP3 (ρ = 0.70, p_adj_ = 5.2×10^−78^), and CD274/PD-L1 (ρ = 0.70, p_adj_ = 1.0×10^−77^). WARS1 (tryptophan-kynurenine axis) and CD80 also ranked among the top associations. ARG1 was the sole gene with a modest negative correlation (ρ = −0.15, p = 4.9×10^−4^).

### 3.3 Dimensionality Reduction Defines IFNG-High and IFNG-Low Transcriptomic Landscapes

PCA of the combined IFN-γ pathway and suppressive TME feature matrix revealed that PC1 (35.1%) and PC2 (9.7%) captured the dominant axes of transcriptomic variation (Figure 5A). IFNG-High tumors (red) clustered towards positive PC1 values while IFNG-Low tumors (blue) occupied negative PC1 space, with partial overlap in the central region consistent with a continuous rather than binary spectrum of IFN-γ activity. The substantial proportion of variance explained by PC1 underscores that IFN-γ pathway activity is the dominant source of transcriptomic heterogeneity in the combined feature space.

**Figure 5.**
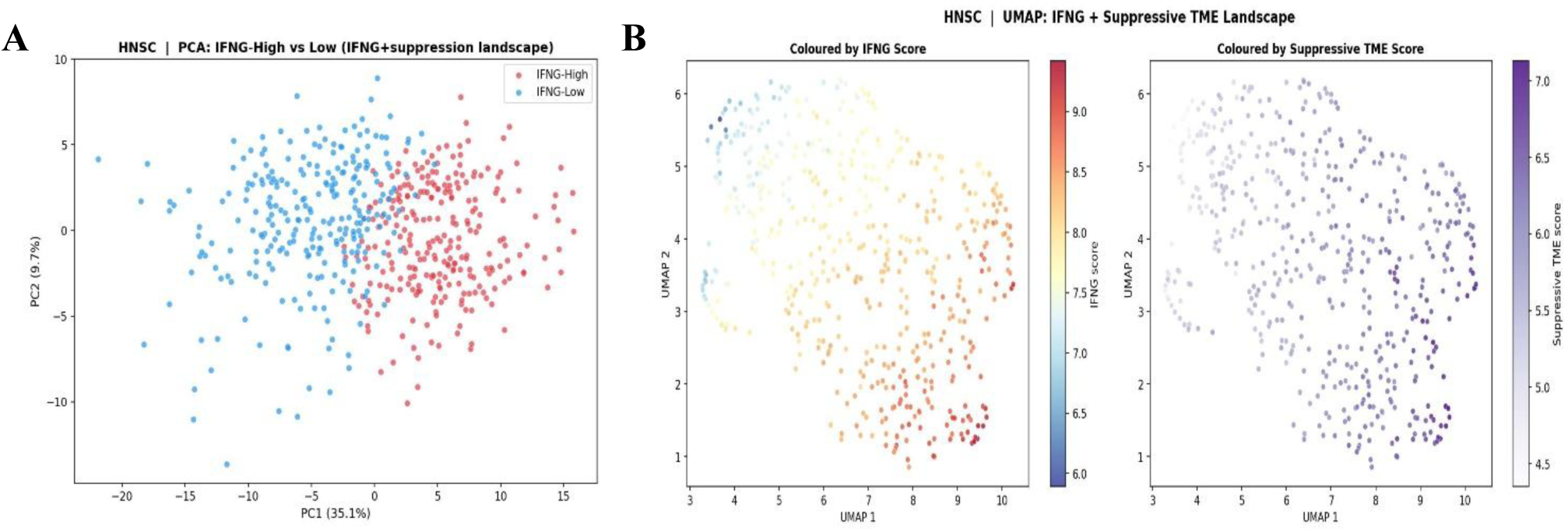
Dimensionality Reduction Reveals IFNG-High and IFNG-Low Transcriptomic Landscapes. (A) PCA scatter plot of HNSC tumors using standardized IFN-γ pathway gene expression and suppressive TME group scores. Points are colored by IFNG-High (red) or IFNG-Low (blue) status (median split of composite IFN-γ score). PC1 (35.1%) and PC2 (9.7%) are labelled. IFNG-High tumors cluster at positive PC1 values. (B) Two-panel UMAP projection of the same feature matrix. Left panel: samples colored by composite IFN-γ score (blue = low, red = high). Right panel: same embedding colored by composite suppressive TME score (white = low, dark purple = high). Co-localization of high IFN-γ and high suppressive TME scores in overlapping UMAP regions confirms coupled immune activation–suppression within individual tumors.

UMAP embedding produced a characteristic crescent-shaped manifold (Figure 5B). Coloring by IFN-γ score revealed a clear cyan-to-red gradient from left to right, while coloring by suppressive TME score showed a complementary white-to-purple gradient in overlapping UMAP regions. Critically, zones of maximal IFN-γ score (lower-right UMAP) co-localized with highest suppressive TME scores, providing a non-linear visual confirmation that immune activation and immunosuppression are co-coupled at the single-tumor level in HNSC.

### 3.4 K-Means Clustering Identifies Four Immunologically Distinct HNSC Subgroups

K-means clustering (k = 4) partitioned HNSC tumors into four immunologically distinct subgroups with clear spatial separation in PCA space (Figure 6A). Cluster 2 (blue) occupied negative PC1 values—corresponding to the IFNG-Low, immunologically cold phenotype. Clusters 3 (orange) and 1 (red) occupied intermediate PC1 positions representing progressive IFN-γ pathway activation. Cluster 4 (green) extended to the most positive PC1 values, indicating the highest IFN-γ pathway and suppressive TME scores.

**Figure 6.**
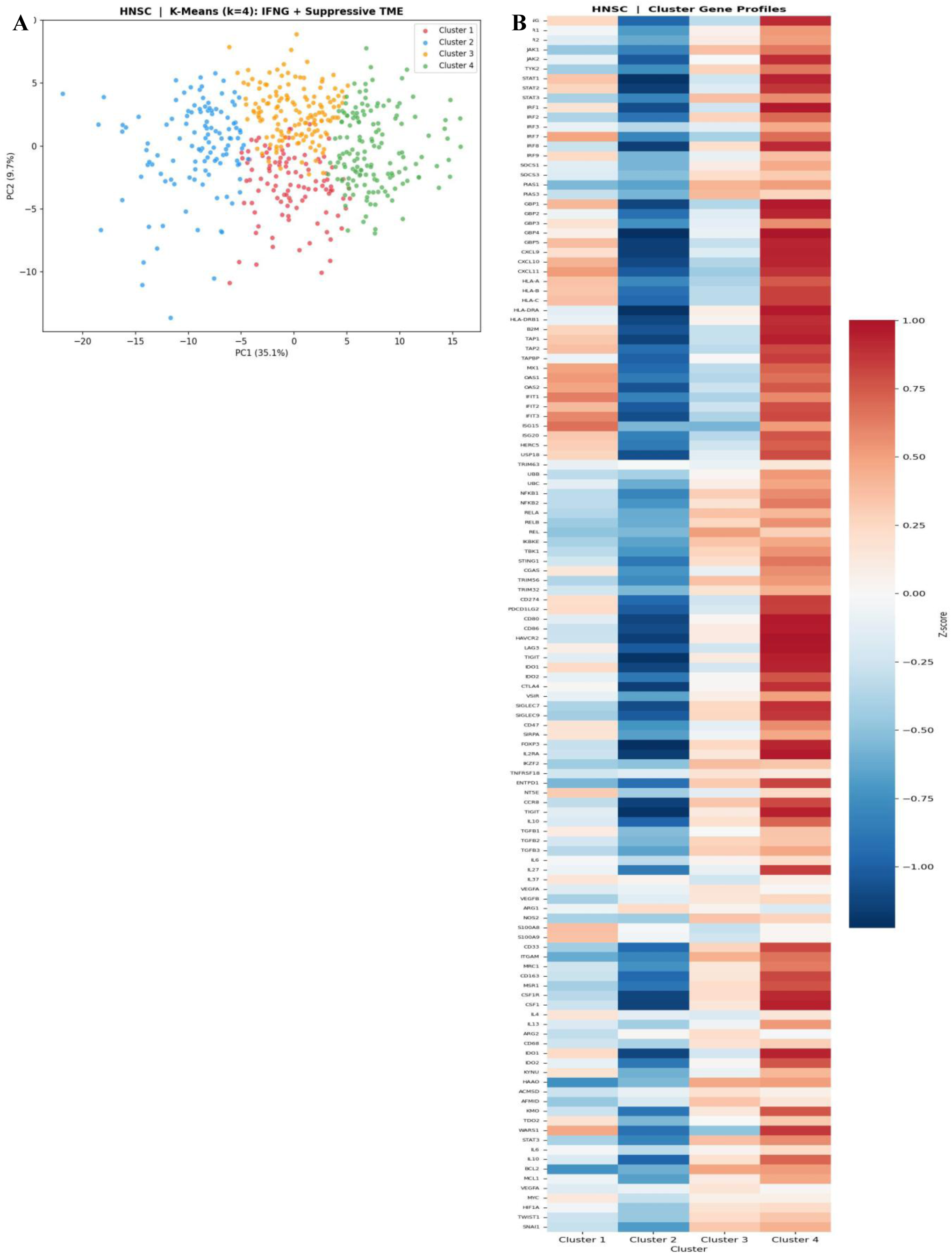
K-Means Clustering (k = 4) Identifies Four Immunologically Distinct HNSC Subgroups. (A) PCA scatter plot (same projection as Figure 3A) with K-means cluster assignments overlaid: Cluster 1 (red), Cluster 2 (blue), Cluster 3 (orange), Cluster 4 (green). The four clusters reveal distinct immunological subtypes with progressive IFN-γ pathway activity along PC1. (B) Z-score-normalized expression heatmap of all IFN-γ pathway and immunosuppressive genes for each cluster (columns: Cluster 1–4). Color scale: deep blue (z < −1) to dark red (z > +1). Cluster 2 is immunologically cold (globally low). Cluster 4 (immune-hot, maximally suppressed) shows simultaneous high expression of IFN-γ effectors and all six immunosuppressive axes, including FOXP3, LAG3, IDO1, and PDCD1LG2.

Cluster gene expression profiles (Figure 6B) revealed the molecular identities of each subgroup. Cluster 2 displayed globally low expression across both the IFN-γ pathway and immunosuppressive genes—an immune-desert phenotype. Cluster 3 showed intermediate IFN-γ activation. Cluster 1 displayed high IFN-γ target gene expression (STAT1, CXCL9/10, GBP1/2, HLA-A) with concurrent checkpoint upregulation. Cluster 4 exhibited the highest expression of both IFN-γ effectors and immunosuppressive genes simultaneously—including FOXP3, LAG3, IDO1, PDCD1LG2, and IL10—defining the immune-hot, maximally suppressed phenotype of primary therapeutic interest.

### 3.5 Random Forest and Permutation Importance Identify IRF8 as the Master Regulator

Random Forest models trained to predict each of six suppressive TME scores from IFN-γ pathway genes revealed distinct feature importance profiles across axes (Figure 1S-A). For Checkpoints_induced, HLA-related and JAK-STAT genes (STAT1, IRF1) dominated. For Treg_induction, STAT1 and HLA-DRA were most important. For MDSCs, TAMs, and Tryptophan/kynurenine, IRF8 emerged as the overwhelmingly dominant predictor (importance ≈ 0.45–0.50), substantially outperforming all other genes. STAT3 and JAK1 led predictions for the STAT3_suppress axis.

Permutation importance analysis on the fitted DNN confirmed IRF8 as the dominant predictor of the composite suppressive TME score by an extraordinary margin (ΔR^2^ ≈ 0.67; Figure 1S-B), more than 15-fold above the second-ranked gene HLA-DRA (ΔR^2^ ≈ 0.04). Secondary contributors included SOCS3 (JAK_STAT), GBP5 (IFNG_targets), and JAK1 (JAK_STAT). The singular dominance of IRF8—a JAK_STAT group member encoding the Interferon Regulatory Factor 8 transcription factor critical for myeloid cell differentiation—identifies it as a transcriptional integrator that amplifies the suppressive arm of IFN-γ signaling, operating in a qualitatively distinct manner from other pathway genes.

### 3.6 DNN Modeling, Network Analysis, and Key Immunosuppressive Correlations

DNN models achieved R^2^ = 0.65–0.85 across six suppressive axes, with Checkpoints_induced (R^2^ = 0.85) and Treg_induction (R^2^ = 0.79) most accurately predicted, and MDSCs and TAMs least (R^2^ = 0.65), suggesting partial IFN-γ independence of myeloid suppression (Figure 2S-A). The DNN architecture diagram (Figure 2S-B) illustrated JAK-STAT genes (orange inputs) as the broadest connectors across output axes. The IFNG pathway–suppression network (Figure 2S-C; |ρ| > 0.35, p < 0.01) revealed a densely positive-correlated hub dominated by JAK-STAT nodes with high degree centrality bridging to all suppressive gene categories.

Scatter plots of the composite IFN-γ score versus 11 key immunosuppressive genes (Figure 2S-D) confirmed the strongest correlations for IDO1 and LAG3 (ρ = 0.80), with TGFB1 more weakly positive (ρ = 0.30) and ARG1 modestly negative (ρ = −0.15). Violin plots (Figure 2S-E) confirmed significant upregulation of all checkpoint and Treg markers in IFNG-High versus IFNG-Low tumors, with the largest median differences observed for IDO1, LAG3, TIGIT, and HAVCR2.

### 3.7 DNN Mediation Analysis Identifies PDCD1LG2 and JAK2 as Key IFNG→PD-L1 Intermediaries

Transcriptome-wide DNN mediation analysis revealed that adding PDCD1LG2 (PD-L2) as a mediator between IFNG and CD274 (PD-L1) produced the largest model improvement (ΔR^2^ = +0.212; total R^2^ = 0.64 versus baseline 0.426; Figure 3S-A). JAK2 ranked second (ΔR^2^ = +0.124; R^2^ = 0.55), followed by GBP5 (ΔR^2^ = +0.094), CD80 (ΔR^2^ = +0.090), CD69 (ΔR^2^ = +0.074), and CXCL11 (ΔR^2^ = +0.074). The mediator landscape bubble plot (Figure 3S-B) placed PDCD1LG2 uniquely at a high indirect score and exceptional DNN gain, while JAK2 combined a high indirect score (∼0.45) with substantial DNN gain, confirming it as both a statistical and mechanistic intermediary.

The DNN architecture diagram (Figure 3S-C) showed the five-mediator model inputting IFNG + PDCD1LG2, JAK2, GBP5, CD80, and CD69 through three hidden layers to predict PD-L1, achieving R^2^ = 0.64 (ΔR^2^ = +0.212 over IFNG alone). The mediator heatmap (Figure 3S-D) confirmed that most top-ranked mediators display strong IFNG-correlated expression gradients, with PDCD1LG2 and JAK2 tracking PD-L1 expression most closely. AC015911.3 and FASLG showed partial inverse patterns, suggesting more complex regulatory inputs.

## 4. DISCUSSION

This study presents the most comprehensive transcriptomic characterization of the IFN-γ immunosuppression paradox in HNSC, employing a multi-modal computational pipeline spanning Spearman correlation analysis, unsupervised dimensionality reduction, K-means clustering, Random Forest regression, deep neural network modelling, permutation importance analysis, JAK-STAT cascade mapping, and DNN-based transcriptome-wide mediation analysis. Collectively, these approaches demonstrate that IFN-γ pathway activity is both the dominant source of transcriptomic variation in HNSC (PC1 = 35.1%) and the primary transcriptional driver of six distinct immunosuppressive axes simultaneously — establishing a quantitative, multi-dimensional portrait of the IFN-γ immunosuppression paradox [28–30].

The universally positive correlations between IFN-γ pathway genes and immunosuppressive axes across all six categories — checkpoint induction, Treg recruitment, anti-inflammatory cytokines, MDSCs/TAMs, tryptophan catabolism, and STAT3-mediated suppression — extend previous observations focused on individual gene pairs. The coexistence of high T-cell recruitment signals (CXCL9/10/11) with high checkpoint expression (PD-L1, LAG3, IDO1) and Treg markers (FOXP3) in the same IFN-γ-high tumors indicates that adaptive immune resistance is not a rare, aberrant response but a fundamental, coordinated counter-regulatory role intrinsic to IFN-γ biology. The ARG1 negative correlation (ρ = −0.15) is the sole exception, suggesting that arginase-mediated MDSC suppression is driven primarily by IL-4/IL-13-polarised myeloid signals in a TME context partially distinct from IFN-γ-dominant inflammation [30–33].

The identification of IRF8 as the dominant transcriptional bridge between IFN-γ signaling and immunosuppression induction (permutation importance ΔR² ≈ 0.67, more than 15-fold above all other genes) is among the most significant findings of this study. IRF8 is a JAK-STAT family transcription factor uniquely positioned at the intersection of IFN-γ signaling and myeloid cell biology [34]. As a direct STAT1 transcriptional target, IRF8 is upregulated by IFN-γ in cells that express it. In the myeloid compartment, IRF8 is a master regulator required for plasmacytoid dendritic cell and cDC1 development, and paradoxically, for MDSC differentiation and function. Loss of IRF8 in tumor-bearing hosts has been reported to reduce MDSC accumulation in preclinical models, and reduced IRF8 expression has been associated with myeloid-biased hematopoiesis that favors immunosuppressive MDSC expansion. In the context of IFN-γ-rich HNSC TMEs, the bimodal IRF8 expression pattern observed in the JAK-STAT cascade heatmap may reflect epigenetically distinct HNSC subpopulations where IRF8 amplifies IFN-γ-induced immunosuppression in a context-dependent manner. Future studies should investigate IRF8 as a therapeutic target for disrupting IFN-γ-to-immunosuppression signal integration [34].

The DNN mediation analysis identified PDCD1LG2 (PD-L2) as the strongest intermediary between IFNG and CD274 (PD-L1) regulation (ΔR² = +0.212). PD-L2 shares the PD-1 receptor with PD-L1 but has received considerably less clinical attention, despite evidence from preclinical and clinical studies suggesting it can independently suppress anti-tumor T-cell activity and mediate resistance to PD-L1-directed therapies. The strong mediation gain of PDCD1LG2 likely reflects shared transcriptional regulation: both CD274 and PDCD1LG2 are direct IRF1/STAT1 target genes with GAS elements in their proximal promoters, creating co-regulated expression from the same IFN-γ-JAK-STAT transcriptional cascade. The variance in PD-L1 prediction captured by PDCD1LG2 but not by IFNG alone may reflect tumor-cell-intrinsic regulatory differences (e.g., amplification of the 9p24.1 locus encoding both PD-L1 and PD-L2, more common in some HNSC subsets) or stromal cell contributions to PD-L2 expression. Therapeutically, this finding reinforces the superiority of PD-1 blockade — which inhibits both PD-L1 and PD-L2 binding to PD-1 — over PD-L1 monotherapy in IFNG-high HNSC [35, 36].

JAK2 as the second-ranked mediator (ΔR² = +0.124) directly implicates the primary IFN-γ signal transduction kinase in determining PD-L1 expression intensity beyond what IFNG level alone predicts. JAK2 expression and activity may serve as an amplifier that scales STAT1/IRF1 transcriptional output per unit of IFN-γ stimulus, with higher JAK2 levels producing disproportionately elevated PD-L1 induction. This suggests that JAK1/2 inhibitors — already FDA-approved for inflammatory conditions and under investigation in oncology — may attenuate IFN-γ-induced PD-L1 upregulation in HNSC. However, JAK inhibition would simultaneously suppress the anti-tumor benefits of IFN-γ signaling (antigen presentation, T-cell recruitment via CXCL9/10/11), necessitating careful evaluation of dose, schedule, and combination approaches [37, 38].

The four K-means clusters provide a clinically meaningful taxonomy of HNSC immune phenotypes [39]. Cluster 2 (immune-cold, globally low IFN-γ and suppressive gene expression) likely corresponds to tumors with low mutational burden, HPV-negativity, and immune exclusion, which are generally least responsive to ICI monotherapy. Cluster 4 — characterized by the co-highest expression of IFN-γ effectors and all six immunosuppressive axes — represents what we term the immune-hot maximally suppressed phenotype. These tumors may contain the most activated tumour-reactive T-cell populations, yet face the greatest immunosuppressive burden. This cluster may be paradoxically the most responsive to ICI combination strategies targeting multiple suppressive axes simultaneously. Clinical correlation of these cluster assignments with ICI response data in existing HNSC trials warrants urgent investigation [40–42].

The DNN models achieved R² = 0.65–0.85 across all six suppressive axes, demonstrating that IFN-γ pathway gene expression is a powerful predictor of immunosuppression magnitude, with Checkpoints_induced most accurately captured (R² = 0.85). This predictive accuracy has practical implications: a composite IFN-γ-based transcriptomic score might serve as a pre-treatment predictor not only of overall immune activation but also of the specific suppressive axes likely to dominate the TME of individual tumors, enabling personalized selection of combination ICI regimens [43, 44].

## 5. CONCLUSIONS

This multi-modal machine learning analysis provides the most comprehensive transcriptomic characterization of the IFN-γ immunosuppression paradox in HNSC to date. We demonstrate that IFN-γ pathway activity is universally and positively correlated with six distinct immunosuppressive axes, confirming that adaptive immune resistance is a fundamental counter-regulatory way intrinsic to IFN-γ biology. Unsupervised clustering identified four immunologically distinct subgroups, including an immune-hot maximally suppressed phenotype most amenable to combination ICI strategies. Deep neural networks predicted suppressive TME scores with R² = 0.65–0.85, establishing a quantitative mechanistic linkage with potential clinical utility. IRF8 emerged as the dominant transcriptional bridge between IFN-γ signaling and immunosuppression (ΔR² ≈ 0.67), while DNN mediation analysis identified PDCD1LG2 (ΔR² = +0.212) and JAK2 (ΔR² = +0.124) as the strongest molecular intermediaries of IFNG-to-PD-L1 regulation, supporting PD-1 blockade superiority over PD-L1 monotherapy. Together, IRF8, PDCD1LG2, and JAK2 constitute prioritized actionable targets for disrupting IFN-γ-driven immunosuppression in HNSC, warranting functional experimental validation and clinical correlation in prospective ICI trials.

## 6. Declarations

## Acknowledgements

Not applicable.

## Author contributions

Eslam E. Abd El-Fattah: Conceptualization, methodology, investigation, writing—original draft preparation, reviewing, and editing. The author read and approved the final manuscript.

Amir Mohamed Abdelhamid: Revision and editing

## Funding

Our article did not receive any financial support from any governmental, private or non-profit organization.

## Code Availability

All code used for data preprocessing, analysis, and figure generation is publicly available at: https://github.com/Eslamabdelfattah/HSNC-IFNG

## Lead contact

Requests for further information and resources should be directed to and will be fulfilled by the lead contact, Eslam E. Abd El-Fattah.

## Materials availability

This study did not generate new unique reagents.

## Availability of data and materials

it is publicly available TCGA data

## Ethics approval and consent to participate

Not applicable.

## Consent for publication

Not applicable.

## Competing interests

We have nothing to declare

## Declaration of generative AI in scientific writing

During the preparation of this work, the author(s) used Copilot in order to improve its readability. After using this tool/service, the author(s) reviewed and edited the content as needed and take(s) full responsibility for the content of the publication

---

**Figure 1S.**
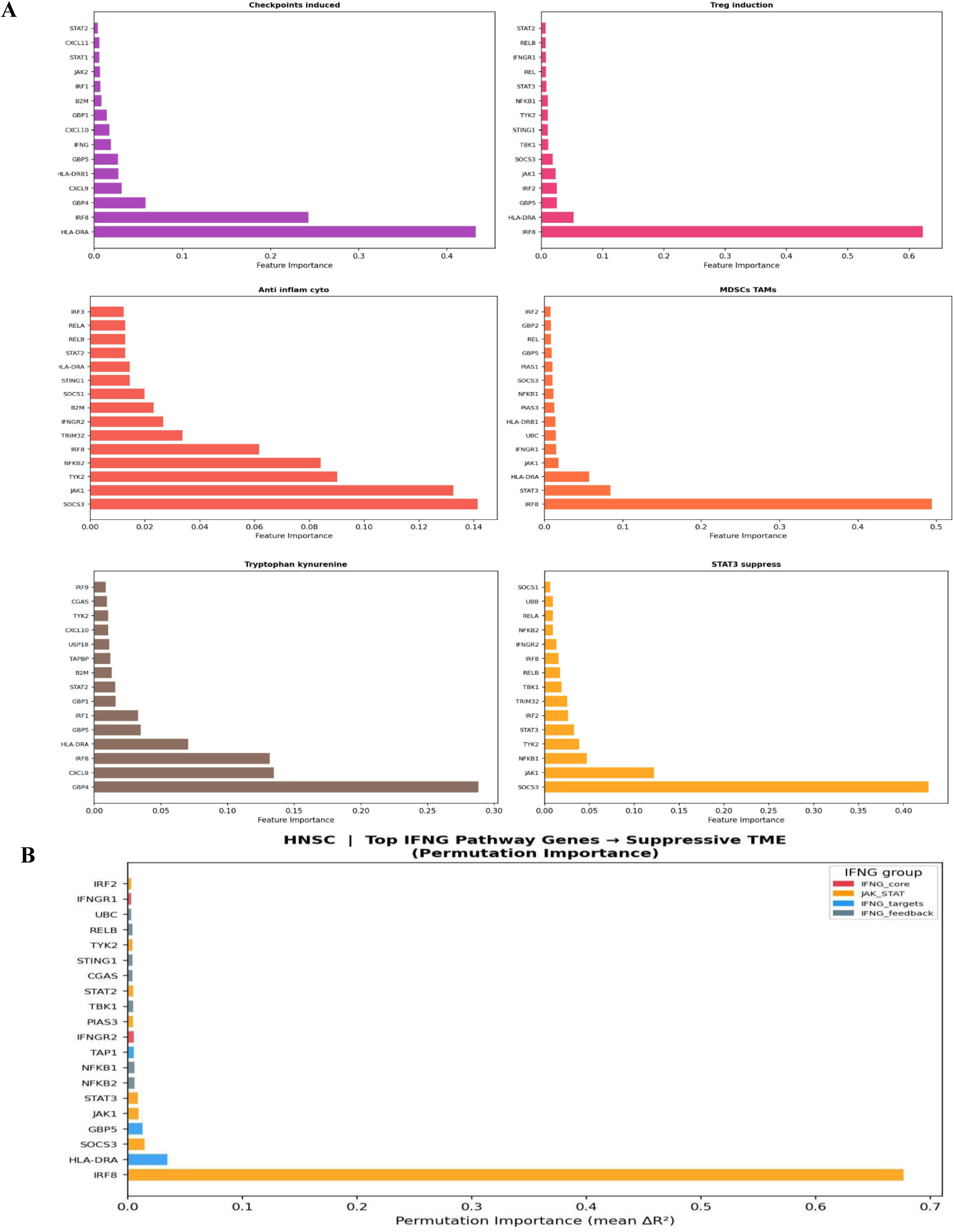
Random Forest Feature Importance and Permutation Importance Identify IRF8 as the Dominant IFN-γ–Suppression Bridge. (A) Random Forest feature importance bar plots (mean impurity decrease) for IFN-γ pathway genes predicting each of six suppressive TME group scores. Bars are color-coded by IFN-γ subgroup: IFNG_core (red), JAK_STAT (orange), IFNG_targets (blue), IFNG_feedback (grey). Each panel represents an independent RF model for one suppressive axis. IRF8 dominates the MDSCs/TAMs and Tryptophan/kynurenine models with importance ≈ 0.45–0.50. (B) Permutation importance (mean ΔR²; x-axis) of IFN-γ pathway genes for predicting the composite suppressive TME score using the fitted DNN model. IRF8 (JAK_STAT; orange) achieves ΔR² ≈ 0.67, more than 15-fold above all other genes. HLA-DRA (ΔR² ≈ 0.04), SOCS3, GBP5, and JAK1 are secondary contributors.

**Figure 2S.**
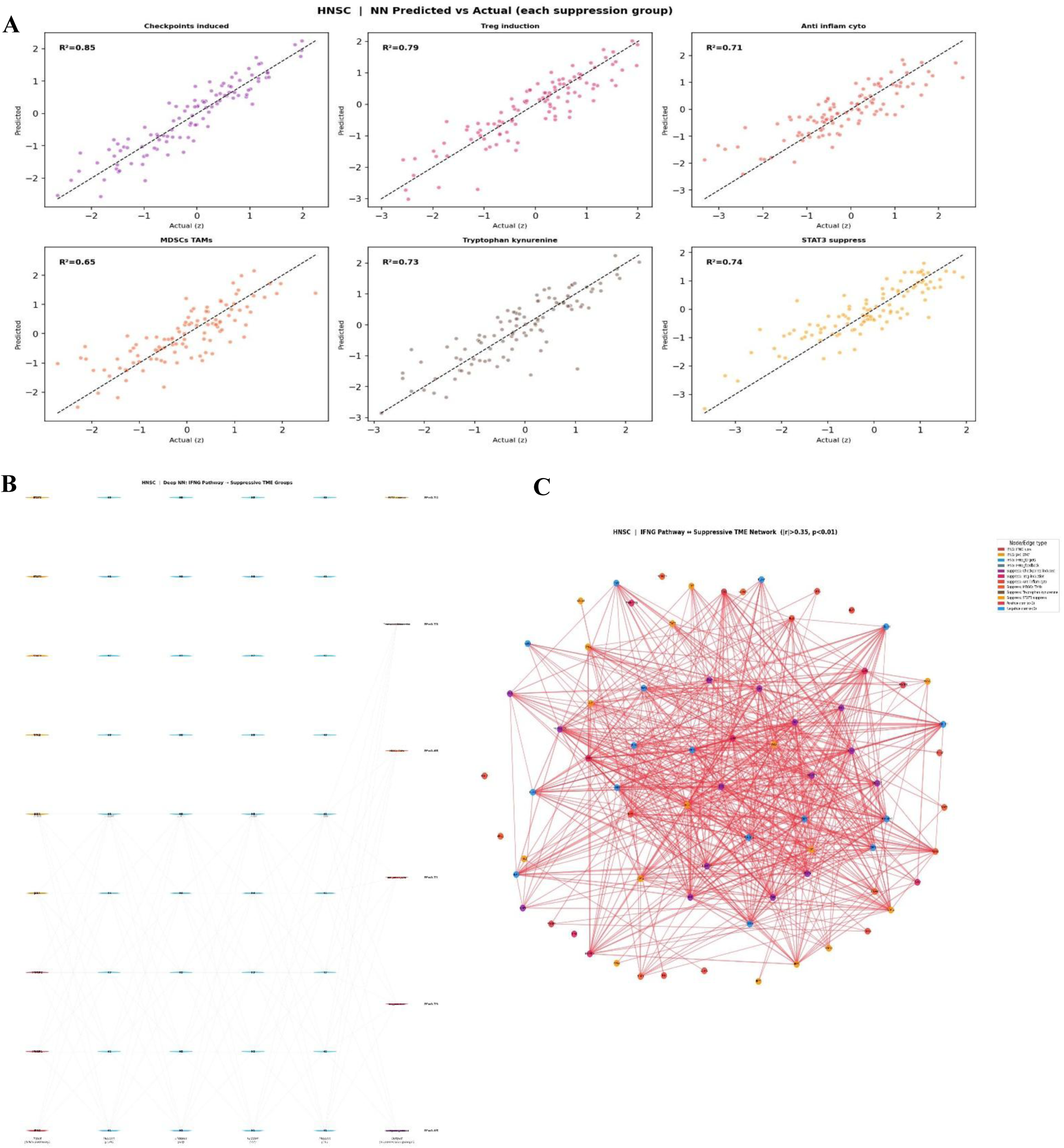
DNN Predictive Models, Correlation Network, and Key IFN-γ–Immunosuppression Relationships. Scatter plots of DNN-predicted versus actual z-normalized suppressive TME group scores (test set) for each of six immunosuppressive axes. R² values are inset; dashed lines indicate perfect prediction. R² ranges from 0.65 (MDSCs/TAMs) to 0.85 (Checkpoints induced). (B) DNN architecture diagram showing input IFN-γ pathway genes (color-coded by subgroup: red = IFNG_score, orange = JAK_STAT, blue = IFNG_targets, grey = IFNG_feedback) through four hidden layers (H1–H4: 128, 64, 32, 16 nodes; cyan) to six suppressive output scores with R² annotations. (C) Correlation network of IFN-γ pathway and immunosuppressive genes (|Spearman ρ| > 0.35, Bonferroni p < 0.01). Node color indicates gene category; edge color: red = positive, blue = negative correlation. JAK-STAT nodes form high-degree central hubs connecting to all suppressive gene clusters. (D) Scatter plots of composite IFN-γ pathway score (x-axis) versus log1p expression of 11 key immunosuppressive genes (y-axis). Spearman ρ and adjusted p-values are annotated. (E) Violin plots comparing log1p expression of the same 11 immunosuppressive genes in IFNG-High (red) versus IFNG-Low (blue) tumors. Spearman ρ and p-values are annotated above each panel.

**Figure 3S.**
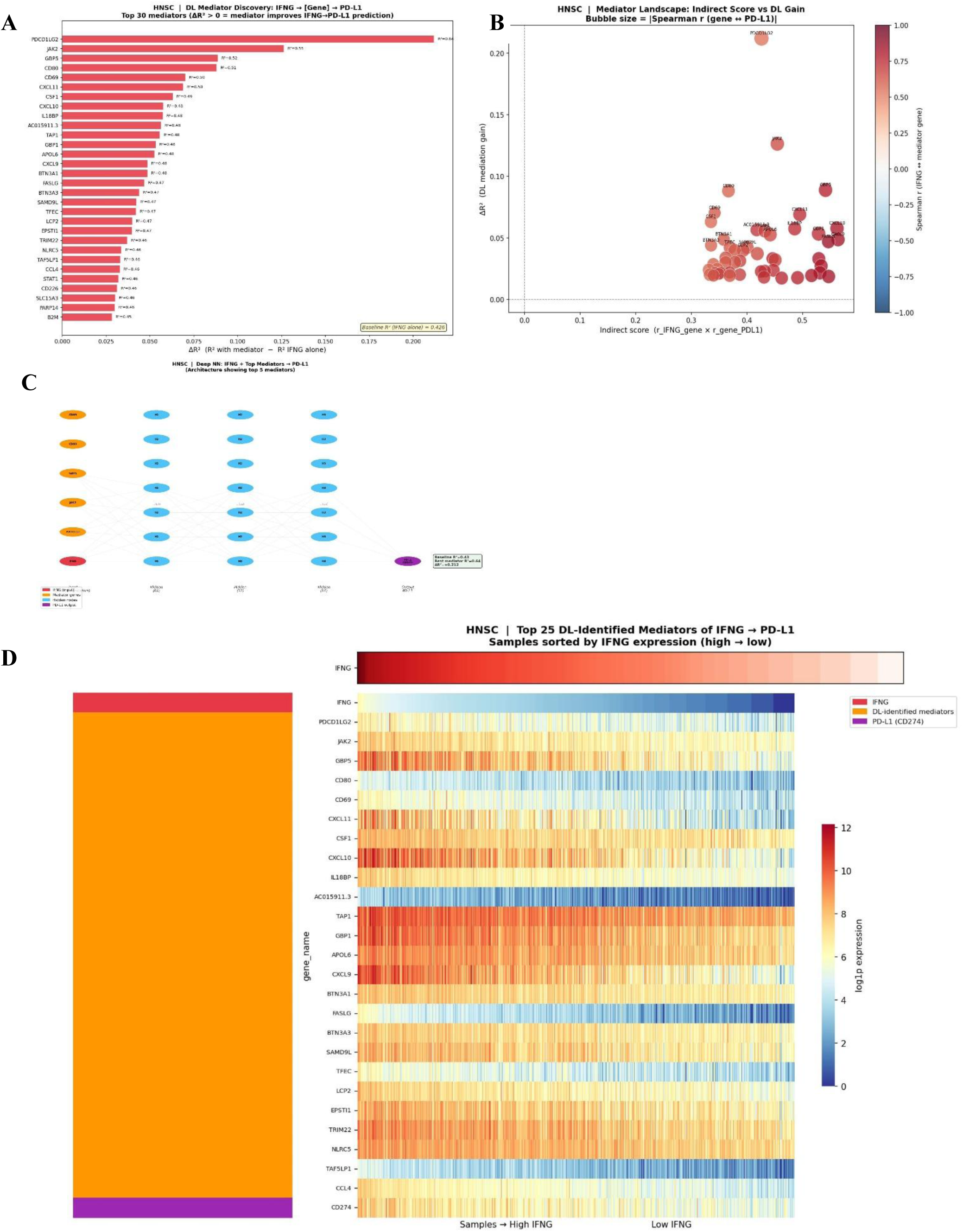
DNN-Based Mediation Analysis: PDCD1LG2 and JAK2 are the Dominant Molecular Intermediaries of IFNG→PD-L1 Signaling. (A) Bar plot of the top 30 mediator genes ranked by ΔR² improvement in DNN prediction of CD274/PD-L1 when each gene is added as a co-predictor with IFNG. R² values with each mediator are annotated. Baseline R² (IFNG alone) = 0.426. PDCD1LG2 (ΔR² = +0.212) and JAK2 (ΔR² = +0.124) are the dominant mediators. (B) Mediator landscape bubble plot: x-axis = indirect effect score [r(IFNG, gene) × r(gene, PD-L1)]; y-axis = DNN mediation gain (ΔR²); bubble size = |Spearman r (gene ↔ PD-L1)|; color = Spearman r (IFNG ↔ mediator gene; red = positive). PDCD1LG2 occupies a unique position with both a high indirect score and an exceptional ΔR². (C) DNN architecture diagram for the top 5 mediator model: IFNG (red) and five mediators — CD69, CD80, GBP5, JAK2, PDCD1LG2 (orange) — as inputs; three hidden layers (64–32–16 nodes; cyan); PD-L1/CD274 output (purple). Annotated: Baseline R² = 0.43, Best mediator R² = 0.64, ΔR² = +0.212. (D) Expression heatmap of the top 25 DNN-identified mediators (orange rows), IFNG (red row), and CD274/PD-L1 (purple row) across HNSC samples sorted by IFNG expression (high → low). Most mediators show graded IFNG-correlated expression, paralleling PD-L1 upregulation.

## Notes

https://github.com/Eslamabdelfattah/HSNC-IFNG

### Competing Interest Statement

The authors have declared no competing interest.

